# Refining structural models of membrane proteins with disordered domains in phospholipid nanodiscs

**DOI:** 10.1101/2022.10.28.512841

**Authors:** Martin Cramer Pedersen, Nicolai Tidemand Johansen, Jennifer Roche, Michael Järvå, Susanna Törnroth-Horsefield, Lise Arleth

**Affiliations:** Niels Bohr Institute, University of Copenhagen, Copenhagen, Denmark; Department of Biochemistry and Structural Biology, Lund University, Lund, Sweden; Department of Biochemistry and Molecular Biology, Gothenburg University, Gothenburg, Sweden

## Abstract

Small-angle scattering can be used to derive structural information about membrane proteins reconstituted in suitable carrier systems enabling solubilization of the membrane proteins in question. Since the studies are done in solution, there is no need for crystallization or deposition on sample grids, and it is in principle possible to obtain structural information about intrinsically disordered regions which cannot be resolved by crystallography or the quantitative link to which is hard to establish using e.g. electron microscopy methods. In this study, tetramers of the gated spinach aquaporin SoPIP2;1 were reconstituted into nanodiscs and small-angle x-ray scattering data were recorded. From these data, we refine structural models of the entire nanodisc-membrane protein complex including the flexible regions using newly developed models based on Fast Debye sums. We introduce software for these computations available via online repositories and discuss the implications and limitations of these methods.

**Author summary:** When it comes to investigating the structure and function of the proteins, a particular class of proteins are known to be cumbersome and problematic: membrane proteins that reside in the cell membrane and regulate and facilitate a number of critical biological processes. Such proteins can often not be studied by conventional means as they unravel and denature structurally or even precipitate in solution. To add insult to injury, such membrane proteins also often contain parts that are intrinsically disordered rendering them irresolvable by e.g. traditional crystallographic techniques and hard to describe structurally. Here, we present a combined computational and experimental approach (as well as the necessary software) to analyze and determine the structure of such proteins in close-to-native conditions in so-called nanodiscs, a biological carrier systems, using small-angle scattering and molecular simulations.

## Introduction

There is an intimate connection between the structure and function of proteins and a detailed understanding of protein function, dynamics, and regulation can be attained via structural studies. By far the most successful method for these types of studies is X-ray crystallography, which is responsible for almost 90% of the close to 185000 structures currently deposited in the Protein Data Bank (PDB) [1] as of October 11th, 2021. While X-ray crystallography may provide structural information at close to atomic resolution, the method is only applicable to proteins that form well-diffracting crystals.

This drawback is particularly noticeable for membrane proteins, whose intrinsic properties make them difficult to crystallize. In particular, the detergent micelle that is commonly used to protect their hydrophobic regions often hinders crystal contacts. As a result, membrane proteins are severely under-represented in the PDB with only 2.5% of the entries describing a membrane protein structure, despite having been estimated to constitute 30% of the human genome [2] as well as 70% of current drug targets [3].

Another limitation of X-ray crystallography is that it is not capable of yielding structural information for proteins or regions of proteins that are flexible or intrinsically disordered. Structural flexibility is an important feature of numerous proteins, allowing them to modulate their function in a dynamic manner. However, such flexibility poses a significant challenge on crystal formation and may hinder the crystallization process altogether. Even if crystals are obtainable, structural information will only be obtained for the parts of the protein with a fixed structure or be limited to conformations that can be confined to the crystal lattice.

Small-angle scattering (SAS) using X-rays (SAXS) or neutrons (SANS) facilitate structural investigations of proteins in solution. Although the obtained structural information is limited to length scales above 10 Å, the technique has the advantage that it circumvents the need for crystallization and allows structural investigtions in more native-like environments. Based on this, small-angle scattering is by now consolidated as a very powerful method for investigating a broad range of soluble proteins and protein complexes [4] and hence a strong complement to high resolution techniques as protein crystallography and nuclear magnetic resonance (NMR). The opportunity for investigating proteins and other particles with molar masses significantly below 100 kDa furthermore allows for covering sizes that are not accessible by Cryogenic Electron microscopy (CryoEM). When it comes to studying membrane proteins by SAS, this is more challenging than for soluble proteins and in recent years, technological and data scientific developments in the CryoEM community have made this technique a key method in the structure determination of membrane proteins. However, analogous to the issues outlined for traditional x-ray crystallography, resolving the three-dimensional structure of flexible domains is also a challenge in CryoEM due to the deposition and/or freezing methods deployed in this technique.

Although in the last few years SAS has improved significantly as a method for obtaining structural information on membrane proteins under solution conditions [5–14], there is still a fundamental need for method development. The membrane protein in question may be investigated while reconstituted in classical storage detergent micelles [5–9, 15] preferably in a size-exclusion SAXS [6, 8, 9] or SANS set up [14, 16], or in more advanced carrier systems such as nanodiscs [10, 12–14]. Nanodiscs are disc-shaped lipoprotein particles consisting of a patch of phospholipid bilayer surrounded by a belt of two copies of a membrane scaffolding protein (MSP) derived from human ApoA1 [10, 17]. This phospholipid bilayer provides a native-like environment which is superior to detergent micelles when it comes to maintaining the functional properties of the incorporated membrane proteins [5]. Furthermore, nanodiscs are homogenous in size and structure [18, 19].

We have previously shown that the monomeric membrane proteins bacteriorhodopsin and proteorhodopsin reconstituted into nanodiscs can be successfully studied using a combination of SAXS and SANS [12, 14]. In this study, we turn our attention to the gated spinach aquaporin SoPIP2;1 as a model system for more complicated membrane proteins with several flexible domains in their structure. SoPIP2;1 is a tetrameric protein with a molecular weight of 120 kDa found in the spinach leaf plasma membrane where each monomer facilitates water transport across the membrane along an osmotic gradient. In response to environmental stress such as drought or flooding, the water permeability of SoPIP2;1 aquaporin is controlled by gating and this has been shown to be triggered by phosphorylation, pH, and binding of Ca^2+^ [20–23].

Crystal structures, an example of which are shown in Fig. 1 (identified by the code 1z98 in PDB), have revealed that this gating mechanism involves a conformational change of a cytoplasmic loop, resulting in the insertion of a leucine residue into the channel opening in the closed state [29]. Neither the N- nor the C-terminus could be fully resolved in the crystal structures of SoPIP2;1, indicating that these regions are intrinsically disordered and highly flexible.

**Fig 1.**
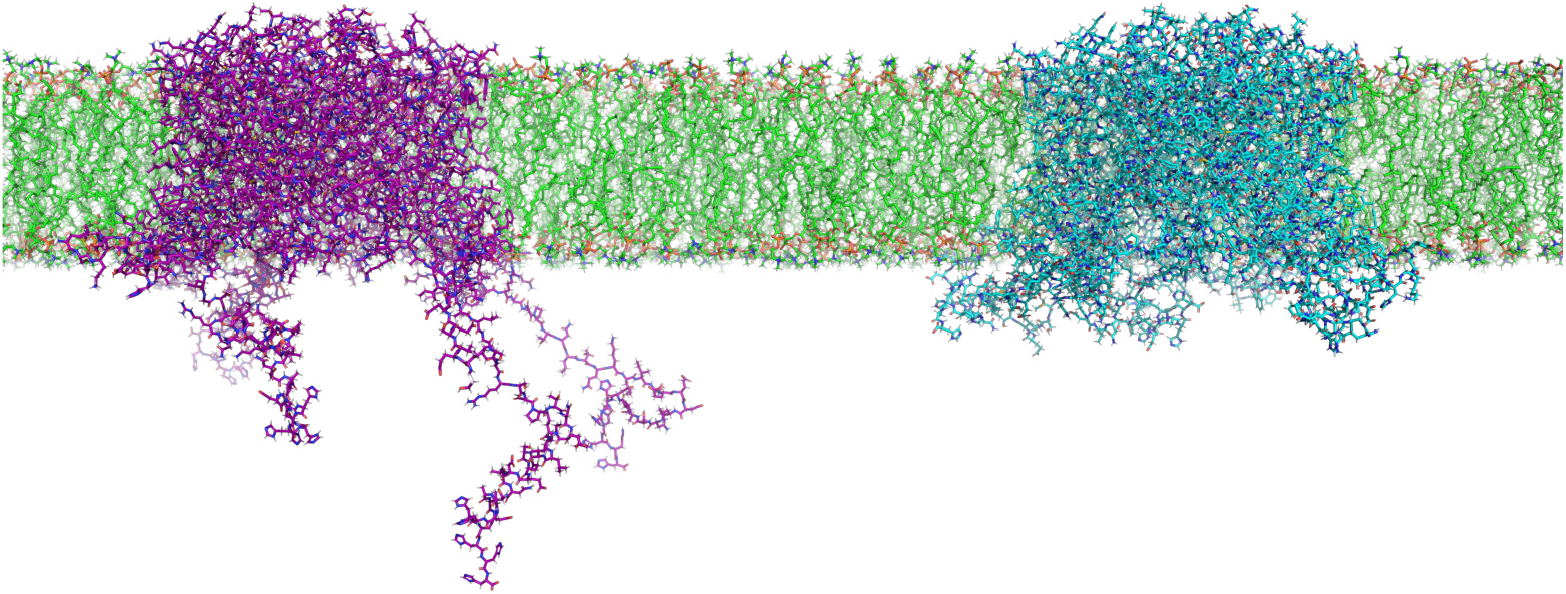
The tetramer 1z98 with extended (on the left, carbon atoms in purple) and contracted (to the right, carbon atoms in cyan) flexible domains embedded in a bilayer of POPC (carbon atoms in green). The structure was assembled with the membrane builder [24–26] in CharmmGUI [27] using the structure from the OPM database [28]. The intracellular, flexible domains were added using the method outlined in this paper.

For this study, SoPIP2;1 was reconstituted in nanodiscs and investigated by SAXS. An optimized sample preparation procedure gave small-angle scattering data of excellent quality, confirming that the structure of SoPIP2;1 inside the disc is in good agreement with the previously determined crystal structure [29]. Using these data, we describe and demonstrate methodological advancements that allows small-angle scattering studies of membrane protein in nanodiscs to be used to gain structural information that could not be obtained by X-ray crystallography.

The problem adressed in this study can be phrased well by considering the amino acid residue sequence of SoPIP2;1 aquaporin, which can be found in Table 1. Simply put, in order to adequately account for the scattering from a sample of SoPIP2;1 aquaporin embedded in a phospholipid nanodisc using the published crystal structure, we must also consider the scattering from the residues missing in the crystal structure.

**Table 1.**
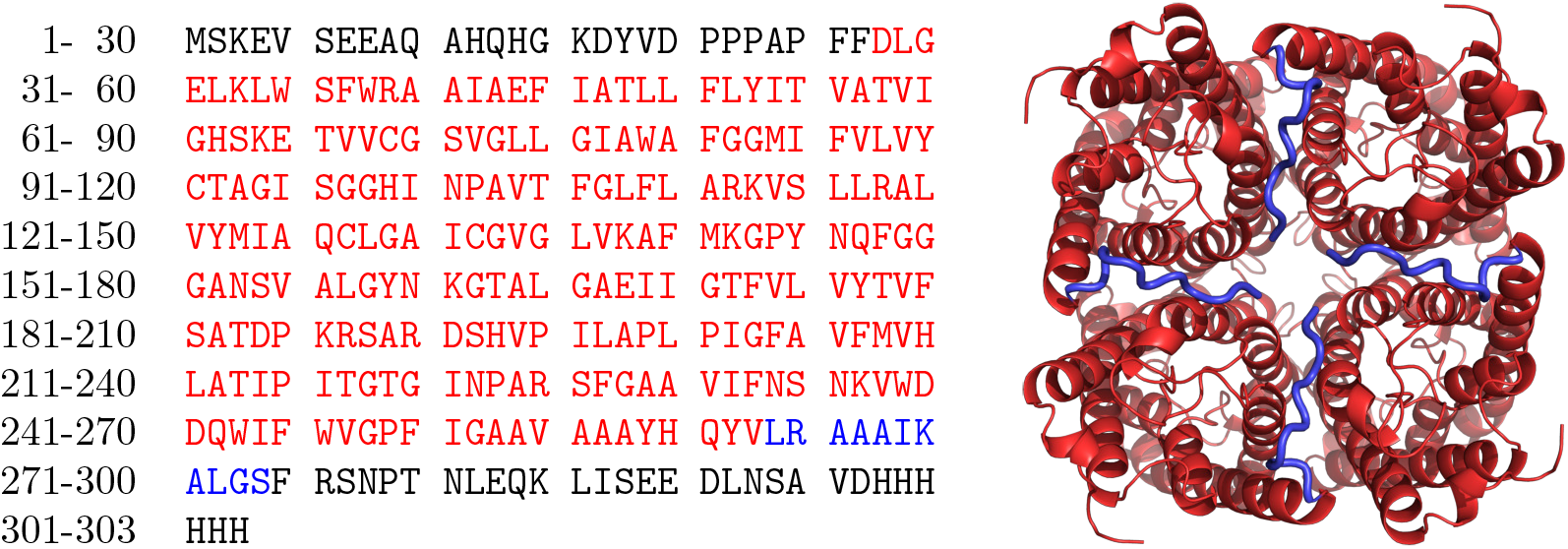
The 303 amino acid residue sequence of SoPIP2;1 aquaporin with the 236 amino acid residues present in the deposited crystal structure 1z98 highlighted in red. Additionally, we consider 11 amino acid residues present in 1z98 in the C-terminal to be flexible in solution based on the crystal structure; these are highlighted in blue (and their cartoon representation emphasized) on the rendering of the 1z98 structure on the right.

|  |  |  |  |  |  |  |
| --- | --- | --- | --- | --- | --- | --- |
| 1- 30 | MSKEV | SEEAQ | AHQHG | KDYVD | PPPAP | FFDLG |
| 31- 60 | ELKLW | SFWRA | AIAEF | IATLL | FLYIT | VATVI |
| 61- 90 | GHSKE | TVVCG | SVGLL | GIAWA | FGGMI | FVLVY |
| 91-120 | CTAGI | SGGHI | NPAVT | FGLFL | ARKVS | LLRAL |
| 121-150 | VYMIA | QCLGA | ICGVG | LVKAF | MKGPY | NQFGG |
| 151-180 | GANSV | ALGYN | KGTAL | GAEII | GTFVL | VYTVF |
| 181-210 | SATDP | KRSAR | DSHVP | ILAPL | PIGFA | VFMVH |
| 211-240 | LATIP | ITGTG | INPAR | SFGAA | VIFNS | NKVWD |
| 241-270 | DQWIF | WVGPF | IGAAV | AAAYH | QYVLR | AAAIK |
| 271-300 | ALGSF | RSNPT | NLEQK | LISEE | DLNSA | VDHHH |
| 301-303 | HHH |  |  |  |  |  |

Inspired by solutions to the similar problem for water-soluble proteins [30, 31], we introduce a pipeline for generating viable structures of the amino acids highlighted in Table 1 using Monte Carlo-based methods. For each monomer consisting of 303 amino acid residues in our tetramer, our models will assume a total of 27 structurally flexible residues in the N-terminal, 236 amino acid residues accounted for in the crystal structure of the aquaporin tetramer, and 40 structurally flexible amino acid residues in the C-terminus. Based on this assumption, we generate an ensemble of conformations of these domains using Monte Carlo-based (MC) methods and use this to analyse our scattering data. Before diving into the technical aspects of the method, we stress that the origin of the structural ensemble of protein conformations is somewhat irrelevant for the outlined methods; specifically, trajectories from molecular dynamics simulations are an obvious source of similar data.

## Materials and Methods

### Cloning, over-expression, and purification of recombinant protein SoPIP2;1

Wild type SoPIP2;1 with a C-terminal Thrombin-site and 6-His-tag was overexpressed in the methylotropic yeast *Pichia pastoris* using 3 L fermentor cultures, typically yielding 280 g cells*/*L culture after 24 h methanol induction. The cells were lysed by a French Press system and the resulting membranes were treated with Urea/NaOH as described previously [32]. The washed membranes were homogenized in resuspension buffer (20 mM Hepes-NaOH, pH 7.0, 50 mM NaCl, 10% w/v glycerol, and 2 mM 2-mercaptoethanol) and stored at -80 °C until further use.

Solubilization was done with drop-wise addition of resuspension buffer with 10% n-octyl-*β*-D-glucoside (Anatrace) to a final concentration of 5%. After incubation at room temperature for 30 min, unsolubilized material was spun down (180000*g*, 30 min, 4 °C) and the supernatant was filtered through a 0.45 µm filter. A Resource S (GE Healthcare) cation exchange column was equilibrated with resuspension buffer with 1% n-octyl-*β*-D-glucoside and the sample was injected and washed with the same buffer. Elution was performed with a linear gradient of NaCl (0 mM to 500 mM) over five column volumes. The peak fractions were collected and concentrated to <500 µL using a VivaSpin concentrator with 10 kDa cut-off (Sartorius Stedim Biotech). The concentrated protein was filtered through a 0.2 µm filter and injected on a Superdex 200 Increase 10/300 GL gel filtration column (GE Healthcare) equilibrated with 20 mM Tris-HCl, pH 7.5, 100 mM NaCl, and 1% n-octyl-*β*-D-glucoside. Peak fractions were pooled and concentrated as described above.

### MSP preparation

MSP1E3D1 was expressed in *E. coli* and purified with slight modifications to a previous described protocol [33]. A culture was grown in 2.5 L TB media at 37 °C in 5 L baffled shaker flasks under agitation at 150 rpm to an OD600 0.6 - 0.8 before induced with ITPG (1 mM) for three hours. The cells were harvested by centrifugation and the resulting pellet was resuspended in 100 mL binding buffer (40 mM TrisHCl, pH 8.0, 300 mM NaCl, 20 mM imidazole) and stored at -20 °C until further processing. The cell suspension was thawed and added Triton X-100 and PMSF to final concentrations of 1% and 1 mM, respectively, before lysis in a cell disruptor (Constant Systems Ltd.). Cell debris was pelleted by centrifugation and the supernatant was loaded onto a self-packed column with NiNTA resin (Qiagen). The column was washed with five column volumes (CV) of binding buffer added 0.1% Triton X-100, five CV of binding buffer added 20 mM Na-cholate, and five CV wash buffer (40 mM TrisHCl, pH 8.0, 300 mM NaCl, 50 mM imidazole). Finally, MSP1E3D1 was eluted in three CV of elution buffer (50 mM Tris-HCl, pH 8.0, 300 mM NaCl, 400 mM imidazole).

The His-tag was removed by enzymatic cleavage, utilising the incorporated TEV-site between the N-terminal His-tag and the MSP1E3D1 sequence. NiNTA purified MSP1E3D1 was mixed with His-tagged TEV protease in a 1 : 100 ratio (w/w TEV:MSP1E3D1) and supplemented with DTT to a final concentration of 1 mM. The mixture was incubated for 20 h at 4 °C while dialysed against TEV buffer (20 mM TrisHCl, pH 7.5, 100 mM NaCl, 1 mM DTT). The self-packed NiNTA column was restored and equilibrated in binding buffer, and non-cleaved MSP1E3D1 and His-tagged TEV protease [34] were removed by flowing the sample over this column. Cleaved MSP1E3D1 was collected in the flow-through fraction and concentrated to 7 mg mL^−1^ in a 10 kDa MWCO centrifugation spin filter (Millipore).

### Nanodisc formation and reconstitution

The reconstitution protocol was based on the procedure developed by the Sligar group [33]. 1-palmitoyl-2-oleoyl-sn-glycero-3-phosphocholine (POPC) was dissolved to a concentration of 50 mM in buffer A (20 mM Tris-HCl, pH 7.5, 100 mM NaCl) containing 100 mM Na-Cholate. The nanodisc mixture was prepared by mixing POPC/Na-Cholate, SoPIP2;1 and MSP1E3D1 to a molar ratio of 210*/*420 : 4 : 2 at a total POPC concentration 10 mM. This mixture was allowed to equilibrate for approximately 2 h at 4 °C with slight agitation.

The detergent was then removed and the nanodisc self-assembled through the addition of 30% w/w Biobeads per sample. After over night incubation, Biobeads were removed by filtration. In order to separate empty and loaded nanodiscs using the His-tag present on SoPIP;1 aquaporin, the sample was injected on a 1 mL HisTrap column (GE Healthcare) equilibrated with Buffer A and washed in Buffer A until the baseline was stable. Loaded nanodiscs were then eluted with Buffer A containing 250 mM Imidazole. Fractions spanning the entire peak area were pooled and concentrated to <500 µL using a 30 kDa cut-off VivaSpin tube (Satorius). This sample was flash frozen and stored at -80 °C until further use.

### Generation of Monte Carlo ensembles

To establish our SAS models, we generated our ensembles of the flexible aquaporin domains using the PHAISTOS software package [35] using the built-in OPLS-MC forcefield. The object is to generate an ensemble of structures capable of reproducing the recorded scattering from the aquaporin-nanodisc complex. The process is charted in Fig. 2.

**Fig 2.**
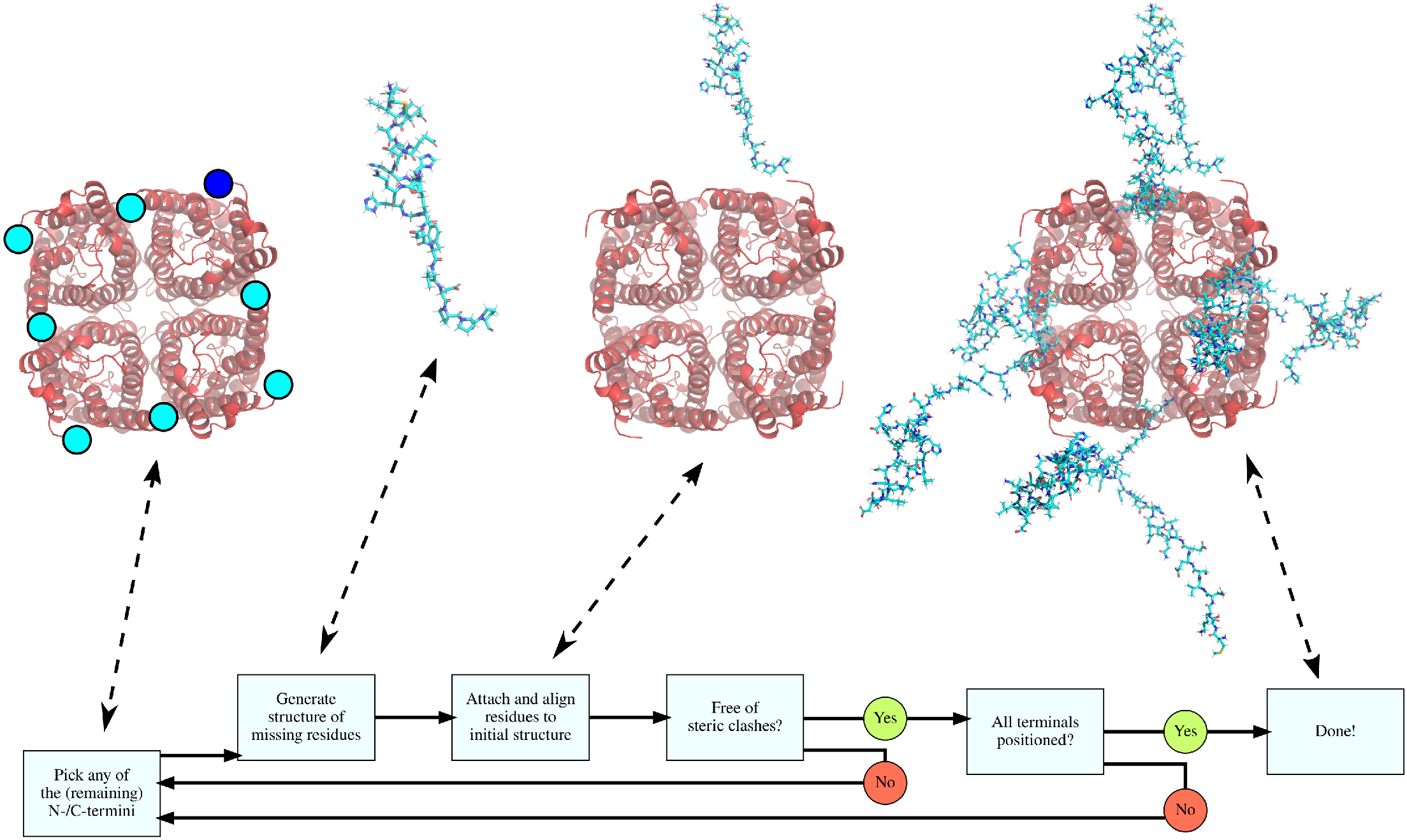
Flowchart describing the process of generating a structure of SoPIP2;1 aquaporin with the amino acid residues not present in the crystal structures added using Monte Carlo ensemble generation. The crystal structure 1z98 (without the amino acid residues highlighted in Table 1) is shown in red and the generated flexible domains are shown in cyan.

Models are discarded if any atom in the generated structure is within the van der Waals radius plus its own van der Waals radius (i.e. a steric clash). Similarly, if an atom is “inside” the hydrophobic part of the eventual bilayer, the generated structure is also discarded.

The handling and input/output of the generated structures were done with the PDB tools [36] for BioPython [37]. With the outlined set up, a viable structure accounting for all 4 × 303 amino acid residues can be generated by PHAISTOS in a matter of seconds on an ordinary laptop.

An initial result from the Monte Carlo ensemble generation became clear after several attempts to generate valid structures of the C-terminal based on the full sequence from 1z98; i.e. with the flexible C-terminal domain beginning after Phe-274. Simply put, we find that the coordinates of the crystal structure are far too constraining to generate 4 non-intersecting models of the C-terminals of 1z98. We kept our model generation scripts running for more than a day with a total of zero structures generated. We conjecture that this reflects the actual physical conditions of the crystallization process and that the coordinates of the C-terminal amino acid residues of 1z98 are to some extent artefacts of the crystallization rather than being representative of the solution structure of SoPIP2;1 aquaporin. Glancing at the visualization of the crystal structure in Table 1, this explanation seems in line with the overall geometry of the tetramer.

For the utilized structure of SoPIP2;1 aquaporin, 1z98, we generated an ensemble of flexible domains not present in the crystal structures. We generated 2000 structures and separated these into two ensembles: the 1000 structures with the most contracted flexible domains and the remaining 1000 with the most extended flexible domains. We determine this by placing the center of the crystal structure at the origin and calculate the moment of inertia. We label our structures as having *contracted* or *extended* flexible domains based on this quantity; i.e. the 1000 structure with the radii of gyration below the median are consider *contracted* and the remaining as *extended*. Fig. 3 is a visualization of this classification. Two examples of the final structures (one with extended flexible domains and one with contracted flexible domains) are shown in Fig. 1 embedded in an computer-based model of a POPC bilayer, and three are shown in Fig. 7, where they have been embedded in the presented nanodisc model.

**Fig 3.**
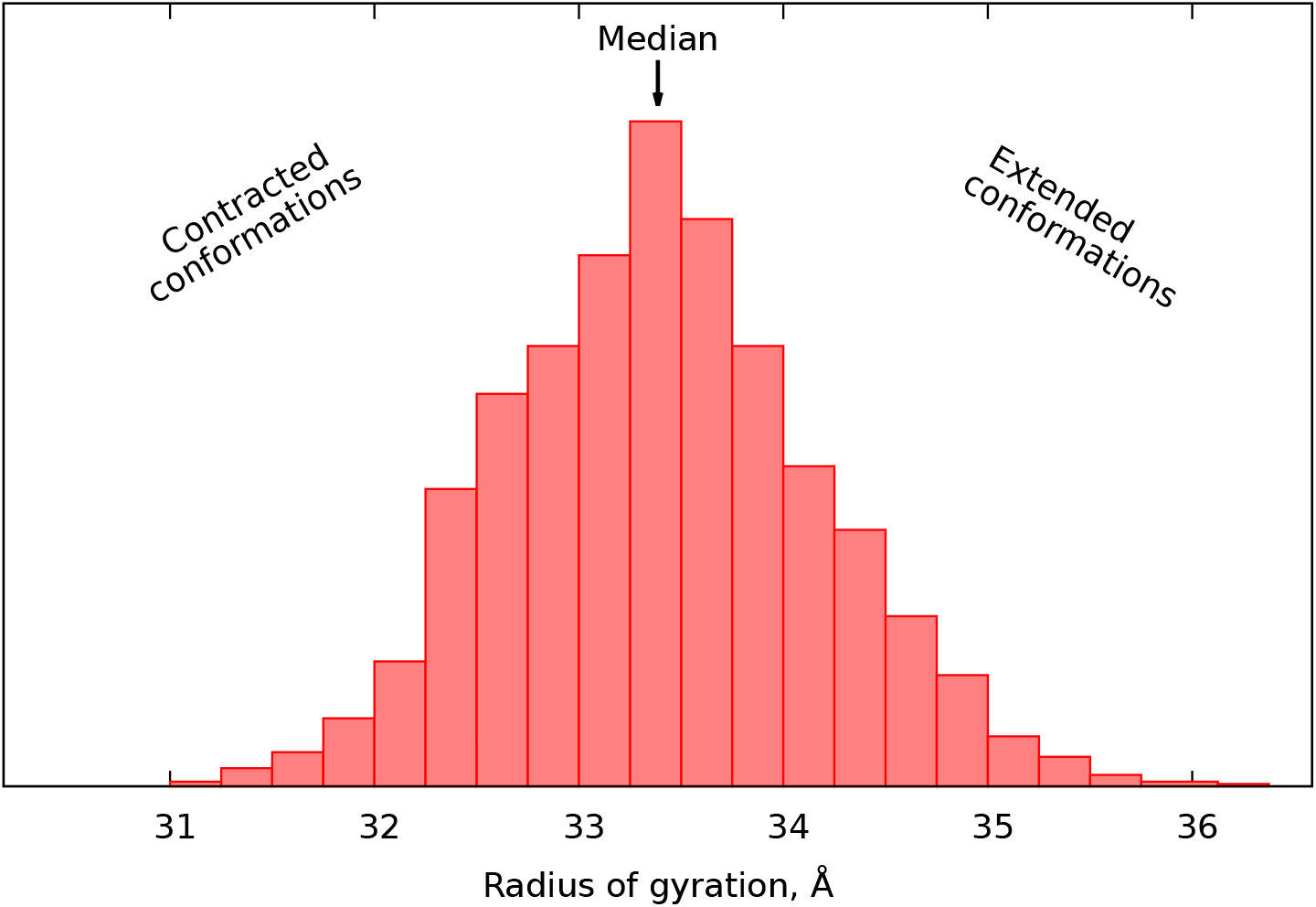
Histogram of the mechanical radii of gyration for our generated ensemble of structures. Structures with radii of gyration below the median are labeled “contracted”, whereas those with radii of gyration above are labeled as “extended”.

### Small-angle scattering

SAXS data were collected at beam line BM29 [38] at the ESRF in Grenoble, France. The data were collected using the SEC-SAXS [39] set up available at this beam line. 180 µL of approximately 8 µM sample was injected onto a Superose6 Increase 10/300 GL column (GE Healthcare) equilibrated in 20 mM TrisHCl pH 7.5, 150 mM NaCl, 10 mM CaCl_2_, 1 mM DTT, and eluted using a flow rate of 0.7 mL min^−1^ with the same buffer. The column was placed at ambient temperature (approximately 20 °C), whereas the SAXS capillary was cooled to 10 °C. One SAXS detector frame per second of exposure was recorded throughout the elution. To improve data statistics, the final SAXS curve was taken as an average of 15 almost identical frames from the right-hand side of the main elution peak as shown in Fig. 4.

**Fig 4.**
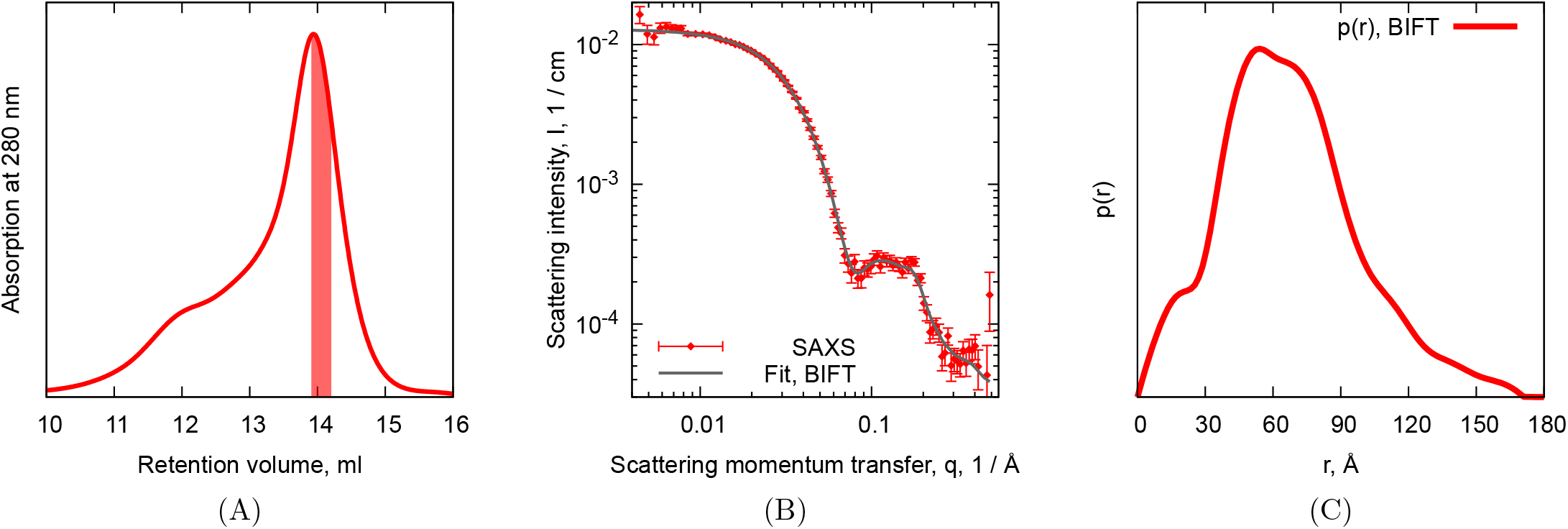
In (A), size-exclusion chromatogram for SoPIP2;1 aquaporin reconstituted into POPC-MSP1E3D1 nanodiscs. The sample was separated on a Superose6 Increase 10/300 column at a flow rate of 0.7 mL min^-1^, while 1 s SAXS frames were collected. SAXS frames from the highlighted area were averaged and used for further analysis. In (B), SAXS data collected on the sample from (A) along with the fits from the Bayesian Indirect Fourier Transform. In (C), the pair distance distribution, *p*(*r*), refined from the data is shown.

We record the scattering intensity, *I*, as a function of the scattering momentum transfer, *q*, which is readily calculated from the incoming wavelength, *λ*, and the scattering angle, *θ*, as *q* = 4*π* sin(*θ*)*/λ*. For the presented SAXS experiments, the nominal wavelengths was 0.99 Å, which with the geometry of the experimental set up allowed us to cover a range of momentum transfer from approximately 0.004 Å^−1^ to 0.5 Å^−1^. The pair distance distribution shown in Fig. 4 was refined using the Bayesian Indirect Fourier Transform algorithm [40, 41] from BayesApp website [42], which is currently avalable at https://genapp.rocks [43] using the default settings for the refinement.

## Results

### SEC-SAXS

Fig. 4(A) shows the size-exclusion chromatography (SEC) profile of SoPIP2;1 aquaporin reconstituted into POPC-MSP1E3D1 nanodiscs. The major peak is observed at the expected retention value for the formed particles. 1 s SAXS frames (corresponding to 0.012 mL of elution volume) were collected throughout sample elution. Due to the presence of a significant shoulder peak, we selected SAXS frames from the right-hand side of the major peak for further analysis (as indicated by the coloured area under the peak). Using the extinction coefficients of our proteins, we estimate the nanodisc concentration in the coloured area to 0.604 µM

Fig. 4(B) shows the averaged SEC-SAXS data. We observe that the overall shape of the data is similar but still somewhat different than that of similar nanodiscs without embedded membrane proteins [14, 44, 45]. The data exhibits the usual Guinier region in the low-*q* part of the data associated with a dilute dispersion of well formed, compact particles. The oscillating features around 0.1 Å^−1^ are usually attributed to the core-shell like structure of a phospholipid bilayer in SAXS, which we expect to see for ordinary nanodiscs. However, the shape is distorted from that usually seen for similar nanodiscs without embedded membrane proteins; as expected for a membrane in which a sizeable membrane protein is embedded.

Accordingly, the refined *p*(*r*) distibution in Fig. 4(C) also exhibits anticipated differences to one refined from data recorded from a sample of nanodiscs without embedded membrane proteins. As an example, the shoulder around 30 Å is considerably more subtle than the local minimum usually observed in data collected from nanodiscs without embedded membrane proteins. This feature is usually attributed to the internal structure of the bilayer (and the negative contrast of the lipid tails); indicating that our membrane protein is seemingly embedded in the bilayer, replacing a significant number of lipids in the bilayer, and thus diminishing this feature. In the process of producing this fit, we obtain a radius of gyration, *R*_*g*_ = 52.3 Å ± 0.1 Å, and a maximum distance between scatterers, *D*_max_ = 169.4 Å ± 2.1 Å, along with a *χ*^2^ of 0.92.

### Modeling and analysis strategy

We begin this section by recapping the Fast Debye Sum approach for calculating small-angle scattering intensities [46, 47]. The scattering *p*(*r*) distribution for a point set of *N* points embedded in 𝔼^3^ in general position, where the *j*th point is equipped with an (excess) scattering length, Δ*b*_*j*_, can be calculated by the double sum:

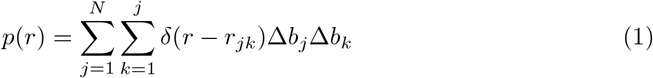

where *r*_*jk*_ is the distance between the *j*th and *k*th point, and *δ*(…) is the Dirac delta function. Consequently, the small-angle scattering intensity can be calculated by Fourier transforming the distribution in Eq. (1):

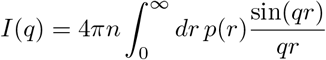

where *n* is the number density. As established in the literature [46, 47], the various components of solution-SAS samples can be described by random point clouds confined to specific regions of space in line with other methods for calculating scattering intensities from geometrical descriptions of compounds [48].

With this mindset, we describe the various components of our nanodisc by randomly generated point clouds confined to geometric shapes specified by the free parameters in the model and the emerging geometry [18, 49]. I.e. rather than calculating the form factor amplitude for e.g. the hydrophobic part of the bilayer, we simply represent it by a dense cloud of points occupying the same space. Our version of the model is parameterized by the following quantities:

- The minor semi-axis of the elliptic cylinder describing the bilayer
- The ratio between the major and the minor axes of the elliptic cylinder describing the bilayer
- The height of the elliptic cylinder describing the hydrophobic part of the bilayer
- The partial specific molecular volume of the lipid, POPC, which implicitly defines the scattering length densities of the lipid bilayer
- The partial specific molecular volume of the membrane scaffolding protein, MSP1E3D1, which similarly defines the scattering length density of the protein belt
- The height of the hollowed-out elliptic cylinder describing the MSP1E3D1-belt, which we fix at 25.78 Å in line with previous studies [45, 50]
- Additionally, an angle describes the in-plane rotation of the aquaporin tetramer relative to the initial orientation of the 1z98 model

Thus, we aim to refine a total of six structural parameters along with a constant background and a parameter accounting for the scale of the model. The specifics and derivations of the geometry of the nanodisc model are discussed elsewhere [18, 49]. A preliminary version of the model included a parameter accounting for the displacement of the aquaporin in the bilayer plane; however, the parameter was refined to close to zero; and consequently, we removed it from the model. For other proteins, adding such a parameter might be a necessity.

As shown in Fig. 5(A), we randomly generate 100000 points inside a box large enough to contain the nanodisc geometry; i.e. (twice the major semi-axis of the bilayer plus the thickness of the MSP belt) × (twice the minor semi-axis of the bilayer plus the thickness of the MSP belt) × (the full height of the bilayer). Depending on the position of each point, we assign the point a scattering length in accordance with the aforementioned geometry; in total, 3 contrast are considered: MSP belt, lipid headgroups, and lipid tails. Points “outside” the nanodisc and points closer than the van der Waals radius to the atoms in the aquaporin structure are discarded; the latter effectively carving out an “aquaporin-shaped” hole through the bilayer of the nanodisc as shown in Fig. 5(B).

**Fig 5.**
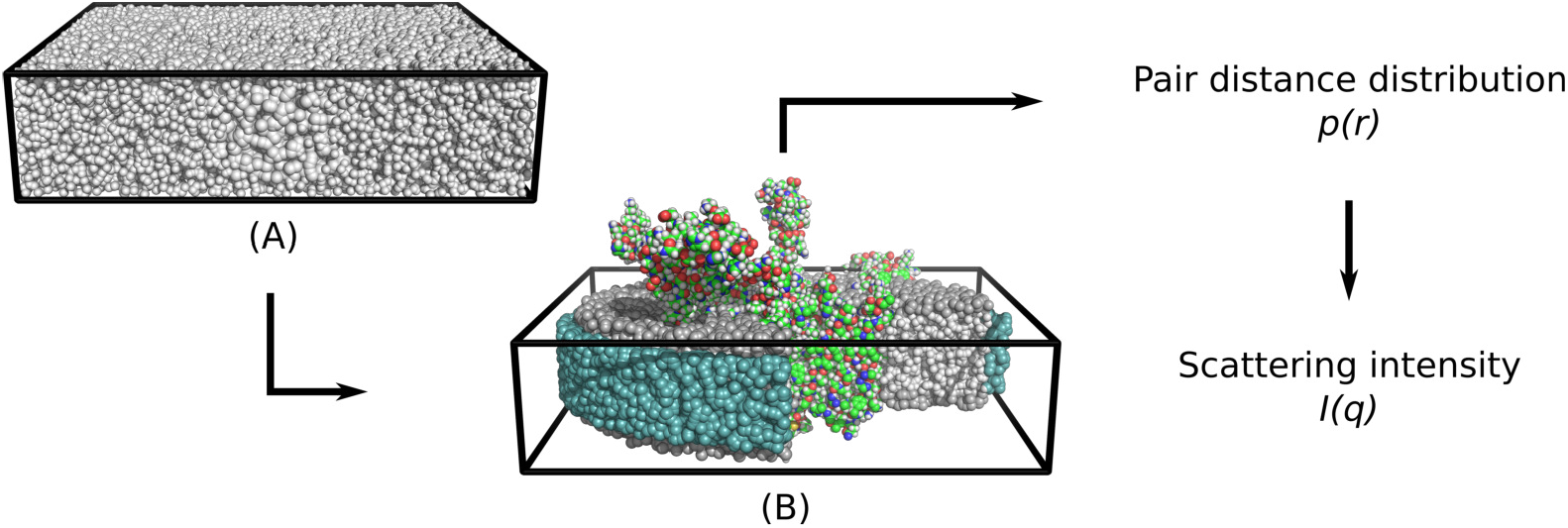
Schematic of the computational pipeline for computing the scattering intensity from a given parameter set describing a nanodisc. In (A), points are randomly generated inside a box of prespecified extent. In (B), these points are either assigned particular scattering properties based on their position to mimick the geometry of the nanodisc or deleted if they i) are too close to the embedded membrane protein or ii) are positioned “outside” the nanodisc. A quarter of the points representing the nanodisc are not shown to emphasize the internal structure of the bilayer.

Key properties of the calculations are sketched in Fig. 6. We see that in the relevant range of momentum transfer, this set up matches the analytical approach in terms of precision. Additionally, we note that 50000 points (or perhaps even 25000) appear to be sufficient when designing models for the nanodisc system in this range of momentum transfer. The presented execution times were calculated using the built-in Linux command time and performed on an ordinary laptop equipped with an NVidia Quadro P2000 GPU. As expected for an algorithm revolving around a double sum, we observe a quadratic-like increase in run times and a typical run time of around 1 s.

**Fig 6.**
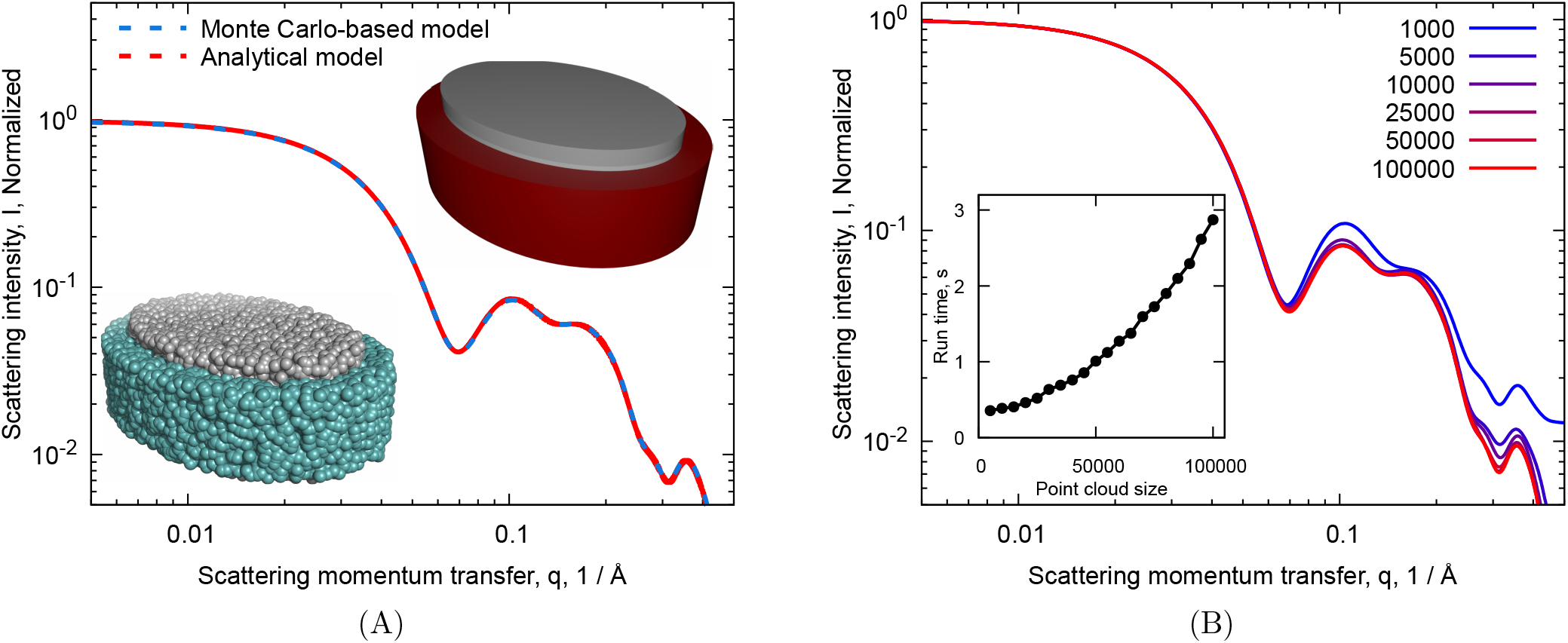
In (A), a proof-of-concept calculation showing the fidelity of the Monte Carlo-based modeling approach when compared to the analytical scattering form factor amplitudes [48] is shown. We calculated the scattering intensity of the well-tested, scattering form factor amplitude-based nanodisc model [18, 49] in WillItFit [51] with default parameters for POPC and MSP1E3D1 and compare it to the same calculation using the Fast Debye Sum-method on a model based on an initial 100000 points. Renderings of the two models are shown with the MSP1E3D1 belt in colors matching the calculated scattering profiles. In (B), we show how the calculated scattering profile changes when the initial number of points is altered; the legend denotes how many points were used in the computation of the nanodisc model. The insert shows the run time of our PyOpenCL-based implementation of the Fast Debye Sum-method versus the size of the input.

The calculations of scattering profiles from our generated models were evaluated in scripts written in OpenCL on an NVidia Titan Xp GPU. These calculations were implemented in a larger refinement framework, WillItFit [51]. We refer the reader to the on-line repository at:

https://gitlab.com/mcpe/FDS.cl for our PyOpenCL-based implementation of the Fast Debye Sum algorithm. This repository also contains example scripts for generating the nanodisc-membrane protein complex models discussed in this manuscript.

Ten examples of scattering intensities calculated from generated aquaporin models embedded in the nanodisc as shown are shown in Fig. 7 to illustrate the variation in scattering generated in this manner. The parameters describing the nanodisc are the ones listed in the left-most column in Table 2.

**Fig 7.**
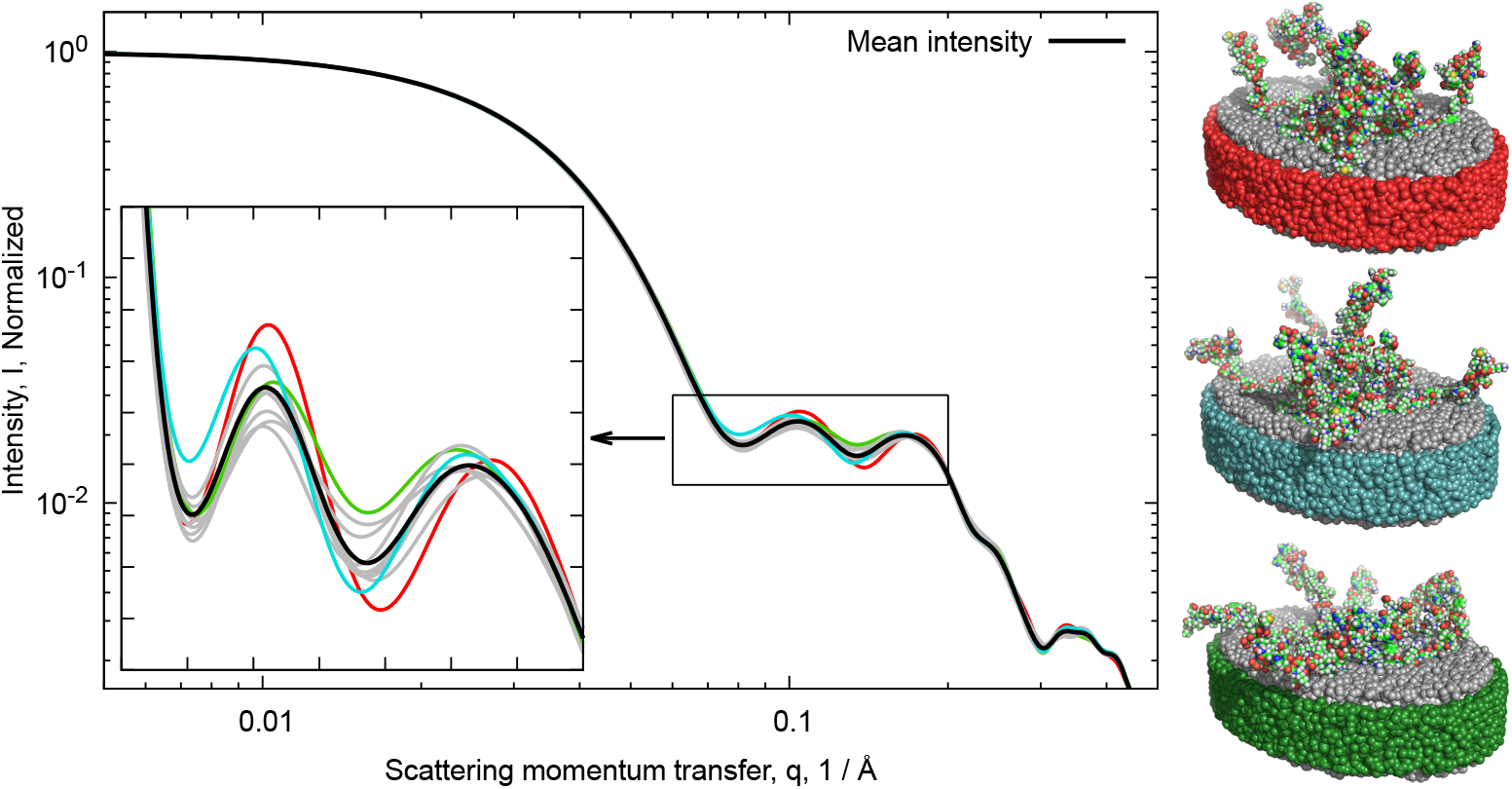
Examples of calculated scattering intensities from ten models in our ensemble with extended flexible domains; the intensities are shown along with the mean scattering intensity of the ten models. Note that the inserted plot has linear axes. Three such models are rendered on the right. The colors of the belt representing the MSP1E3D1 proteins are chosen to match the corresponding curves of the left.

**Table 2.** On top, parameters refined from the data shown in Fig. 8 using the presented models. Below, selected parameters calculated from the upper ones. The error estimates represent 68.3% confidence intervals [52, 53].

| Refined parameters | All structures | Extended domains | Contracted domains |
| --- | --- | --- | --- |
| Axis ratio of bilayer | $1.41 \pm 0.372$ | $1.41 \pm 0.463$ | $1.40 \pm 0.390$ |
| Minor semi-axis of bilayer, Å | $51.6 \pm 10.9$ | $51.5 \pm 12.6$ | $51.9 \pm 12.6$ |
| Hydrophobic height of bilayer, Å | $29.5 \pm 6.57$ | $29.8 \pm 5.29$ | $29.4 \pm 5.97$ |
| Volume of POPC, Å<sup>3</sup> | $1266 \pm 123$ | $1261 \pm 109$ | $1268 \pm 115$ |
| Volume of MSP1E3D1, Å<sup>3</sup> | $33000 \pm 11000$ | $33600 \pm 11600$ | $32400 \pm 11800$ |
| Angle of rotation of aquaporin, rad | $1.21 \pm 1.09$ | $1.05 \pm 0.270$ | $1.01 \pm 0.614$ |
| Scale | $0.774 \pm 0.250$ | $0.775 \pm 0.243$ | $0.754 \pm 0.212$ |
| Background, $10^{-5} \text{ cm}^{-1}$ | $2.61 \pm 3.40$ | $2.76 \pm 3.47$ | $2.67 \pm 3.29$ |
| **Other parameters** |  |  |  |
| $\chi^2$ | 1.15 | 1.17 | 1.17 |
| Major semi-axis of bilayer, Å | 72.9 | 72.5 | 72.5 |
| Thickness of MSP1E3D1 belt, Å | 6.56 | 6.36 | 6.12 |
| Density of MSP1E3D1 belt, $\text{g cm}^{-3}$ | 1.64 | 1.61 | 1.67 |
| Height of bilayer, Å | 39.6 | 40.0 | 39.6 |
| Area per lipid headgroup, Å<sup>2</sup> | 64.0 | 63.0 | 64.1 |
| Lipids per nanodisc | 232 | 233 | 232 |
| Lipids displaced by aquaporin | 137 | 139 | 137 |

We conclude this section by considering the computational implications of the presented Fast Debye Sum-approach to the problem of calculating scattering intensities. First of all, we note that the parallelization via GPUs is computationally attractive though similar GPU-based schemes might be employable in the context of form factor-based models. To our knowledge, this has yet to be attempted. The choice of number of points used to represent a given shape depends entirely on the desired range in momentum transfer, for which the subsequent scattering intensity profile is to be calculated. The higher the upper bound on *q*, the higher the resolution and the more points are needed to describe a continuous structure.

Secondly, in this set up, the time-consuming calculation of pair distance distributions from our point cloud is independent from the number of datapoints, for which the intensity is to be calculated. For the presented values, the subsequent Fourier transformation is quite negligible in terms of computational resources. In terms of scaling behavior, we note that the construction of the *p*(*r*) distribution is quadratic in number of points due to the double sum in Eq. 1.

Calculating the scattering intensity profile from a single conformation on the right in Fig. 7 takes a few seconds on a modern GPU. While this is seemingly rapid and tractable, it does imply that evaluating the scattering profiles for an entire set of 1000 conformations of the flexible domains of aquaporin does take an hour or two. Consequently, utilising this model inside an optimization routine to refine structural parameters from small-angle scattering data is a task that might take several days.

As another boon, modeling complex biophysical systems such as the ones presented here makes preliminary design, coding, and debugging a simpler task, as the models are constructed in real space rather than reciprocal space as is often the case for such systems. The generated point clouds are readily inspected for trivial errors through visualizations of the computed model. This is a considerably harder task for e.g. form factor amplitude-based models for scattering. Such practical advantages of this style of modeling makes data analysis an easier task for a community wider than the consolidated scattering users.

### Model refinement

We refine three distinct structural models from the presented scattering data: based on either the 1000 most *extended* conformations, the 1000 most *contracted* conformations, or the full set of 2000 conformations for the flexible domains. The corresponding fits are shown in Fig. 8. The parameters refined in the fitting process are listed in Table 2. We discuss these further in the coming section.

**Fig 8.**
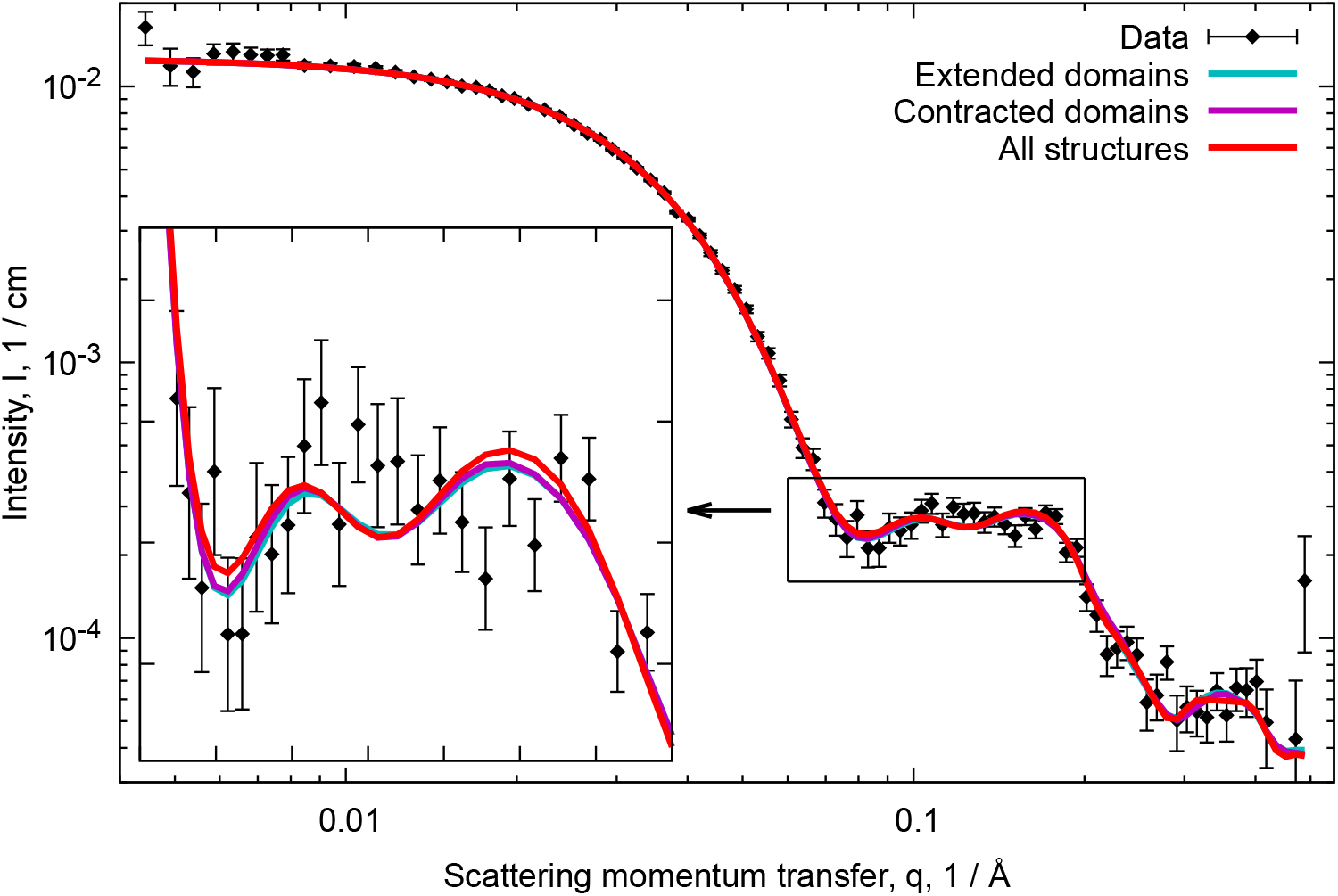
Fits of the models based on *extended* flexible domains, *contracted* flexible domains, or the full set of both types of structures. Note that the insert has linear axes.

We find that the presented models are all capable of reproducing the general features in the data as evidenced by the higher *χ*^2^. In particular, we note good agreement in the Guinier (i.e. low-*q*) region of the data, implying that the model captures the overall shape of the irradiated complex well. The models capture the descending slope in the low-*q* region of the data as well as the bumps around *q* = 0.12 Å^−1^ and *q* = 0.18 Å^−1^.

## Discussion

The refined partial specific molecular volume of POPC (see Table 2) is in good agreement with reported values [54, 55]. Similarly, we observe that the area per phospholipid headgroup for POPC is close to the traditionally quoted value from the literature [55] as well as published values for similar nanodiscs without embedded membrane proteins [44, 45]. Comparable results have been refined before from similar systems [12, 14] implying that the embedded membrane protein might induce slight perturbation in the structure of the surrounding lipid bilayer.

Perhaps unsurprisingly, our model is unable to firmly ascertain certain parameters in our models. In particular, we note the large error estimates on the orientation of the aquaporin tetramer in the bilayer and on the axes of the bilayer patch. The first is perhaps to be expected due to the symmetry of the nanodisc and aquaporin tetramer. We explain the latter by considering how the flexible nature of the sample might obfusciate the information on the dimensions of nanodisc in the scattering signal. Similarly, we note how our estimates of the volume of MSP1E3D1 are much more uncertain than in reports where only the nanodisc itself was investigated. As the MSP accounts for only about a third of the protein content in the sample, this is perhaps to be expected. This does to some extent mitigate the concerns in the previous paragraph. In fact, the refined volume of the MSP1E3D1 belts surrounding the bilayer patch is to some extent in accordance with previously published values [14, 44]. We note that as reported for other membrane scaffolding proteins, MSP1E3D1 appears to be denser than the average protein, to which the literature usually associates a density of 1.35 g cm^−3^ [56]. Consequently, the width of the protein belt is somewhat lower than what one might expect. This is in line with previous reports on several members from this family of membrane scaffolding proteins [44, 45]. However, we do stress the uncertainty associated with this value.

The point cloud-based SAS models utilized in the study differ fundamentally from the form factor-based models often utilized in this field that we and others have been promoting until now. As a consequence of this, one technically obtain different values of *χ*^2^ when rerunning a model computation due to slight differences in the randomly generated point clouds. Effectively, this gives us a manner in which to estimate whether we are generating sufficiently many points for a model refined from a given dataset. These differences in *χ*^2^ must be negligible when performing two model computations with the same parameters. We refer the reader to Fig. 6 for a demonstration of the validity of our modeling choices. On the topic of *χ*^2^, we note *en passant* that all three models produce *χ*^2^s above that of the BIFT algorithm, implying that the models are as such not overfitted considering the apparent noise level of the presented data [57].

One of the main benefits of the outlined approach is the manner in which it is readily extended to molecular dynamics trajectories of membrane proteins. In this study, we based our scattering model on structures generated by Monte Carlo-based sampling of protein conformations using PHAISTOS, but sampling these from any trajectory sweeping the relevant conformational space of the given protein is entirely feasible. As molecular dynamics often constrain simulations using information from e.g. NMR experiments, we believe this to be a promising approach in the field of integrative structural biology. In this context, it is worth noting that from a computational perspective, MC is usually considerably less demanding than MD in terms of sweeping a conformational configuration space of a protein structures. Naturally, the physics of the system in question is taken into account in entirely different manners in the two approaches. Small-angle neutron scattering (SANS) is another source of data, which is readily utilized and incorporated in the outlined approach. The models used in this study are trivially generalized to the neutron contrast; a method which is frequently used in scattering physics of biophysical systems.

In terms of post-analysis of trajectories or Monte Carlo-generated ensembles, we believe the outlined method could synergize well with e.g. the Ensemble Optimization Method (EOM) [30] or Bayesian Reweighing methods [58, 59] for refining narrower conformational ensembles from the initial pool of structures.

## Conclusions

In this paper, we presented a new modeling strategy for analysing and refining structural models from small-angle scattering data from membrane proteins with disordered domains embedded in phospholipid bilayer nanodiscs. Monte Carlo-based simulations proved a powerful tool for producing the conformational ensemble needed to construct viable models for application in small-angle scattering model refinement; though we stress that other sources such as MD or NMR will be just as viable.

We discussed the computational implications of the presented modeling strategy, which we believe to be favorable for a number of systems where the computational burden of handling conformational ensembles is too intense using other methods. Our code is available on-line.

Aquaporin served as an ideal model system for the presented study and method development despite the recorded data on the system not allowing us to distinguish between the various conformational ensembles. We conjecture that the outlined approach is particularly well suited for membrane proteins with large, flexible domains [60, 61] as these will contribute relatively much and comparably more to the scattering signal.

With the recent developments in synchrotron technology and construction of new, powerful neutron science and synchrotron facilities as well as the continouos evolution of beam lines at these facilities, the limits for which samples can be investigated using SANS and SAXS are constantly pushed. Additionally, advanced sample handling and sample environment set ups allow for probing the structure of more elaborate and dynamic biophysical samples [14, 16, 39].

## Acknowledgements

The authors gratefully acknowledge SAXS beamtime at the BioSAXS Beamline, BM29 at ESRF Grenoble. We thank Dr. Petra Pernot from the ESRF for the help to facilitate optimal execution of the experiments, and Abigail Barclay for her aid in testing the presented software. NVidia is acknowledged for granting an NVidia Titan Xp GPU kindly provided via the NVidia Academic Seeding Grants. Furthermore, we thank the Novo Nordisk Foundation for funding this project via the Synergy programme, grant number NNF15OC0016670, and the Lundbeck Foundation for similarly financing this study via the Brainstruc programme, grant number R155-2015-2666.

